# Phosphatidylcholine biosynthesis in Mitis group streptococci via host metabolite scavenging

**DOI:** 10.1101/664672

**Authors:** Luke R. Joyce, Ziqiang Guan, Kelli L. Palmer

**Affiliations:** Department of Biological Sciences, The University of Texas at Dallas, Richardson, Texas, USA; Department of Biochemistry, Duke University Medical Center, Durham, NC

**Author notes:** Corresponding authors: Ziqiang Guan, Kelli Palmer.

## Abstract

The Mitis group streptococci include the major human pathogen *Streptococcus pneumoniae* and the opportunistic pathogens *S. mitis* and *S. oralis* which are human oral cavity colonizers and agents of bacteremia and infective endocarditis in immunocompromised patients. Bacterial membrane lipids play crucial roles in microbe-host interactions, yet for many pathogens, the composition of the membrane is poorly understood. In this study, we characterized the lipidomes of selected species of Mitis group streptococci and investigated the mechanistic basis for biosynthesis of the phospholipid phosphatidylcholine (PC). PC is a major lipid in eukaryotic cellular membranes, but it is considered to be comparatively rare in bacterial taxa. Using liquid chromatography/mass spectrometry (LC/MS) in conjunction with stable isotope tracing, we determined that Mitis group streptococci synthesize PC via the rare host metabolite scavenging pathway, the glycerophosphocholine (GPC) pathway, which is largely uncharacterized in bacteria. Our work demonstrates that Mitis group streptococci including *S. pneumoniae* remodel their membrane in response to the major human metabolites GPC and lysoPC.

**Importance:** We lack fundamental information about the composition of the cellular membrane even for the best studied pathogens of critical significance for human health. The Mitis group streptococci are closely linked to humans in health and disease, yet their membrane biology is poorly understood. Here, we demonstrate that these streptococci scavenge major human metabolites and use them to synthesize the membrane phospholipid phosphatidylcholine. Our work is significant because it identifies a mechanism by which the major human pathogen *S. pneumoniae* and the primary human oral colonizers *S. mitis* and *S. oralis* remodel their membrane in response to host metabolites.

## Introduction

The Mitis group streptococci are Gram-positive bacteria that natively inhabit the human oral cavity, nasopharynx, and gastrointestinal tract (1). They include the species *S. mitis* and *S. oralis*, which are among the first colonizers of the human oral cavity from birth and facilitate host-microbe-microbe interactions by creating anchors for biofilm formation with other oral microbiota (2, 3). *S. mitis* and *S. oralis* are also opportunistic pathogens causing bacteremia and infective endocarditis (4–7). The Mitis group streptococci also include the major human pathogen, *S. pneumoniae*. *S. pneumoniae* has >99% 16S rRNA sequence identity (8–10), exchanges capsule biosynthesis and antibiotic resistance genes (11, 12), and shows antibody cross-reactivity (13) with *S. mitis* and *S. oralis*.

We, and others, recently reported that certain Mitis group streptococci have unusual membrane physiology in that they can proliferate while lacking the major anionic phospholipids phosphatidylglycerol (PG) and cardiolipin (CL) (14, 15). More specifically, *S. mitis* and *S. oralis* can tolerate deletion or inactivation of *cdsA*, the gene encoding phosphatidate cytidylyltransferase (14, 15). CdsA catalyzes synthesis of cytidine diphosphate-diacylglycerol (CDP-DAG), a key intermediate in the synthesis of major phospholipids (Fig 1). Deletion or inactivation of *cdsA* in Mitis group streptococci, and the corresponding loss of PG and CL, confers high-level (MIC >256 µg/mL) resistance to daptomycin, a last-line lipopeptide antibiotic (14–16).

**Fig 1.**
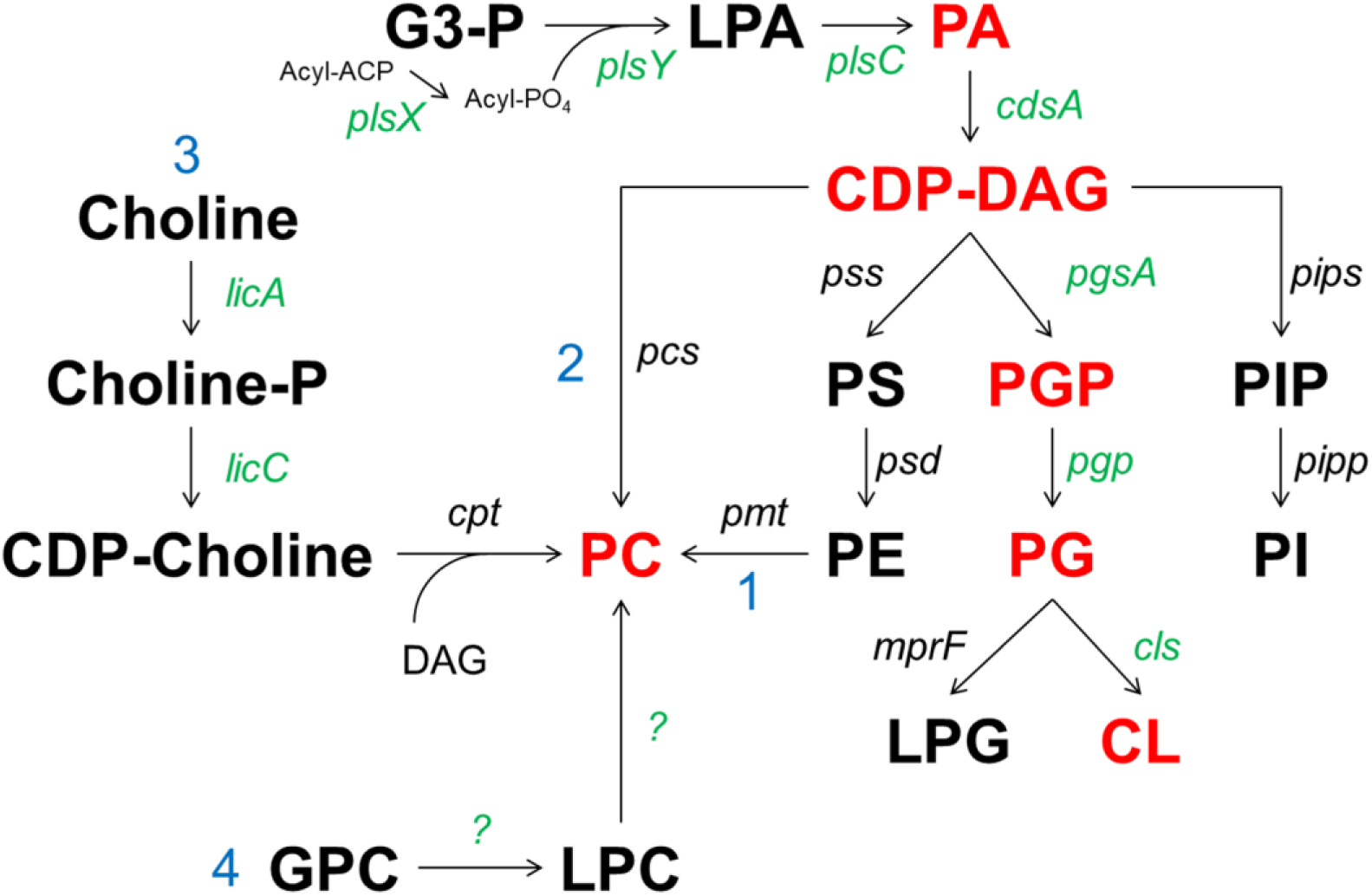
Phospholipid pathways in bacteria, including pathways for PC biosynthesis. Shown are the methylation pathway (1), phosphatidylcholine synthase pathway (2), CDP-choline pathway (3), and the glycerophosphocholine pathway (4). Lipids detected in THB-cultured SM43 are shown in red. Genes encoded by SM43 are shown in green. Abbreviations are PA, phosphatidic acid; CDP-DAG, CDP-diacylglycerol; PS, phosphatidylserine; PE, phosphatidylethanolamine; G3-P, glycerol-3-phosphate; PGP, phosphatidylglycerol-3-P; PG, phosphatidylglycerol; CL, cardiolipin; LPG, lysly-phosphatidylglycerol; PIP, phosphatidylinositolphosphate; PI, phosphatidylinosotiol; PC, phosphatidylcholine, LPC, lysophosphatidylcholine; GPC, glycerophosphocholine; LPA, lysophosphatidic acid.

Unexpectedly, lipidomic analysis of *S. mitis* and *S. oralis* by normal phase liquid chromatography-electrospray ionization/mass spectrometry (NPLC-ESI/MS) revealed the presence of phosphatidylcholine (PC) in wild-type strains and in *cdsA*-null mutants (14). To our knowledge, PC in streptococci had not been previously described. Overall, the lipid compositions of streptococci are under-studied and poorly characterized. Previous studies analyzing lipids of streptococci primarily used thin layer chromatography, whose limitations in analytical sensitivity and molecular specificity prohibit comprehensive lipidomic identification; these studies did not detect PC (17–22). PC is a biologically significant lipid. As a zwitterionic phospholipid, PC promotes bilayer formation (23), reduces the rate of protein folding to allow for correct protein configurations (24, 25), aids in resistance to antimicrobials targeting prokaryotic membranes (26), aids in surviving environmental fluctuations such as temperature shifts (27), and is a major component of eukaryotic membranes. There is evidence that PC plays important roles in host-microbe interactions. *Legionella* lacking functional PC biosynthesis exhibit decreased virulence because of poor recognition by host macrophages and reduced motility (28). *Brucella abortus* and *Agrobacterium tumefaciens* also exhibit diminished virulence when PC biosynthesis is inactivated (29–31). Alternatively, *Pseudomonas aeruginosa* strains lacking PC show no detectable alteration in virulence (32).

Because PC may impact how Mitis group streptococci interact with the human host, in this study we investigated the mechanism for PC biosynthesis in these organisms. There are four experimentally confirmed PC biosynthesis pathways in bacteria (Fig 1), two of which are widespread and well-characterized, namely the phosphatidylethanolamine (PE) methylation pathway (23, 33), and the phosphatidylcholine synthase (Pcs) pathway (34, 35). The PE methylation pathway uses PE as a starting substrate, which is methylated via phospholipid *N*-methyltransferase (Pmt) in three subsequent steps to form monomethylPE (MMPE), dimethylPE (DMPE), and finally PC, using *S*-adenosylmethionine as the methyl group donor (33). The PE methylation pathway is utilized by mammalian liver cells (23, 36) and bacteria including *Rhodobacter sphaeroides* and *Sinorhizobium meliloti* (23). The Pcs pathway is exclusive to prokaryotes and is a one-step reaction in which a phosphatidylcholine synthase (Pcs) enzyme condenses choline with CDP-DAG to form PC (34). The presence of either *pmt* or *pcs* genes has been used to identify bacterial taxa likely to produce PC (23).

A third pathway, the CDP-choline pathway (referred to as the Kennedy pathway in eukaryotes), was recently identified in the Gram-negative human oral colonizer, *Treponema denticola* (37). In this pathway, choline is scavenged from the environment and activated to CDP-choline via the LicAC enzymes. Many host-associated bacteria possess LicAC and utilize host-derived choline to decorate a wide range of extracellular structures (23) including the Type IV lipoteichoic acid (LTA) of *S. pneumoniae, S. mitis*, and *S. oralis* (38–42). In the CDP-choline pathway, CDP-choline is condensed with diacylglycerol (DAG) by a 1,2-diacylglycerol choline phosphotransferase (CPT) to form PC.

A fourth pathway, the glycerophosphocholine (GPC) pathway, has been reported for only two organisms: the Gram-negative plant pathogen *Xanthomonas campestris* (43) and yeast (44). In eukaryotic cells, GPC is a breakdown product of choline-containing membrane phospholipids. Yeast can utilize GPC as the source for glycerol-3-phosphate, choline, or phosphate, depending on the environmental conditions (45). GPC is a major human metabolite present in saliva and blood (46, 47). In the GPC pathway, GPC is scavenged from the environment and acylated twice to form the intermediate lysoPC and then PC. The genetics underlying the GPC pathway in *X. campestris* have not been fully elucidated. Moser *et al*. identified two *X. campestris* acyltransferases that performed the second acylation event from lysoPC to PC (43). Yeast possess a fully elucidated GPC pathway (44, 48).

Here we use NPLC-ESI/MS and other biochemical and genetic approaches to investigate PC biosynthesis in Mitis group streptococci using type strains and an infective endocarditis isolate (Table 1). We determined that these organisms synthesize PC by the rare GPC pathway via scavenging the host metabolites GPC and lysoPC.

**Table 1.** Summary of major glycolipids and phospholipids detected in Mitis group streptococci in routine laboratory media^1^.

| Streptococcal species | Strain | Source | DAG | MHDAG | DHDAG | PA | PG | CL | C55-P | PC | L-PG | Growth medium<sup>2</sup> |
| --- | --- | --- | --- | --- | --- | --- | --- | --- | --- | --- | --- | --- |
| *S. mitis* | ATCC 49456 | Type strain | + | + | + | + | + | + | + | + | - | THB |
| *S. oralis* | ATCC 35037 | Type strain | + | + | + | + | + | + | + | + | - | THB |
|  | 1647<sup>3</sup> | Endocarditis | + | + | + | + | + | + | + | + | - | THB |
|  | 1648<sup>3</sup> | Endocarditis | + | + | + | + | + | + | + | + | - | THB |
| *S. pneumoniae* | D39 | Historical strain | + | + | + | + | + | + | + | + | - | THB+Y |
|  | TIGR4 | Bacteremia | + | + | + | + | + | + | + | + | - | THB+Y |
| Mitis group | 1643 (SM43)<sup>3</sup> | Endocarditis | + | + | + | + | + | + | + | + | - | THB |
|  | SM43Δ*cdsA* | This study | + | + | + | + | - | - | + | + | - | THB |
|  | SM43Δ*pgsA* | This study | + | + | + | + | - | - | + | + | - | THB |

<sup>1</sup>Abbreviations: DAG, diacylglycerol; MHDAG, monohexosyldiacylglycerol; DHDAG, dihexosyldiacylglycerol; PA, phosphatidic acid; PG, phosphatidylglycerol; CL, cardiolipin; C55-P, undecaprenyl phosphate; PC, phosphatidylcholine; L-PG, lysyl-phosphatidylglycerol
<sup>2</sup>Todd-Hewitt Broth (THB); THB supplemented with 0.5% yeast extract (THB+Y).
<sup>3</sup>Lipid profiles previously published by Adams *et al* 2017 (14).

## Results

### PC biosynthesis by model *S. mitis* and *S. oralis* strains used in this study

We previously reported lipidomic analysis by NPLC-ESI/MS of three Mitis group infective endocarditis isolates cultured in the rich, undefined laboratory growth medium Todd Hewitt broth (THB). We determined that these organisms possess PC in their membranes ((14) and Table 1). We confirmed these results using the *S. mitis* and *S. oralis* type strains ATCC 49456 and ATCC 35037, respectively (Table 1).

In this study, we used one of the infective endocarditis isolates, referred to as SM43, for most of our mechanistic studies of PC biosynthesis. Phylogenetic assignments within the Mitis group are difficult due to variable phenotypes and highly conserved 16S rRNA sequences (8, 9). SM43 was initially assigned to the *S. mitis* species using standard biochemical techniques (49). We analyzed a complete SM43 genome sequence using an average nucleotide identity (ANI) calculator (50, 51). ANI values are used for molecular species definitions (52). Two bacterial strains with an ANI of ≥95% in their shared genes are considered to be the same species and with an ANI of >70% are considered to be the same genus (53, 54). SM43 possesses 94.3% ANI with *S. oralis* ATCC 35037 and 86.9% ANI with *S. mitis* ATCC 49456. Based on these data showing a close phylogenetic relationship between SM43 and *S. oralis*, we refer to SM43 as a Mitis group *Streptococcus* in this study (Table 1).

### PC biosynthesis in SM43 is not by the PE methylation or Pcs pathway

The well-characterized PE methylation pathway for PC biosynthesis is catalyzed by the enzyme Pmt (Fig 1). This pathway is excluded as SM43 does not synthesize PE (14), nor does it encode *pmt*.

The Pcs pathway requires CDP-DAG and choline as substrates (Fig 1). *cdsA* loss-of-function mutants arising spontaneously under daptomycin selection in Mitis group streptococci do not synthesize CDP-DAG but still synthesize PC (14). These results indicate that PC biosynthesis in Mitis group streptococci is by a CDP-DAG-independent pathway. To confirm these results, we generated a *cdsA* deletion in SM43. PC was detected in both wild-type and Δ*cdsA* SM43 cultured in THB. Fig 2a shows the relative abundance of PC in the SM43 membrane during mid-exponential phase. The ESI/MS spectra of the major PC species, PC (30:1) at *m/z* 704, PC (32:1) at *m/z* 732, PC (34:1) at *m/z* 760, and PC (36:2) at *m/z* 786 are shown for SM43 (Fig 2b) and *S. mitis* type strain ATCC 49456 (Fig S1a). The chemical structure of PC (34:1) shown in Fig 2c is supported by MS/MS (Fig S1b). The ESI/MS spectra of PC in SM43Δ*cdsA* is shown in Fig 2d. Based on these results and the absence of a *pcs* ortholog in the SM43 and ATCC 49456 genomes (Fig 1), the *pcs* pathway is excluded.

**Fig 2.**
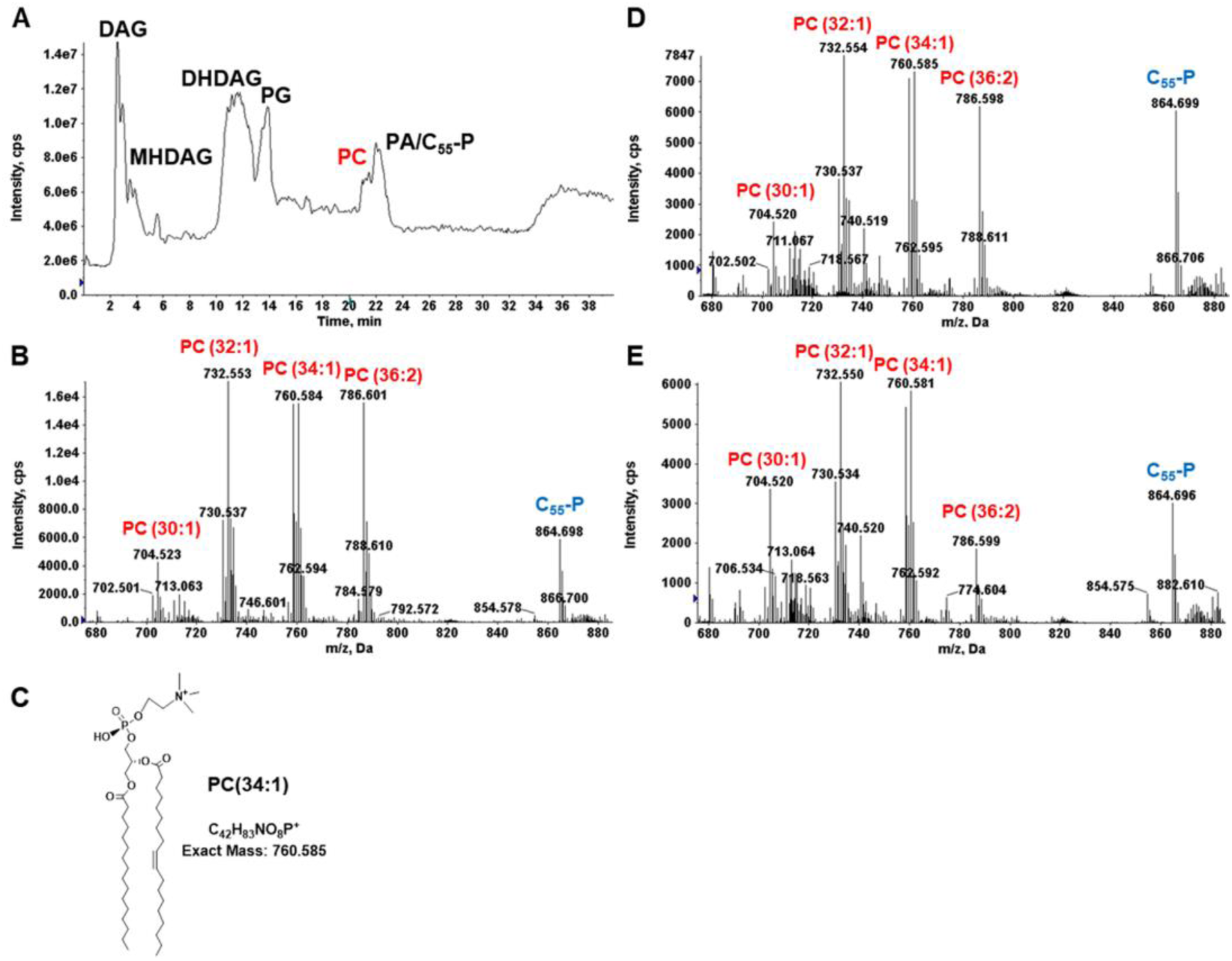
PC species detected in SM43, SM43Δ*cdsA* and SM43Δ*pgsA*. Shown are the positive mode total ion chromatogram of SM43 lipids (A), the ESI-MS of SM43 PC species (B), chemical structure of PC (34:1) (C), the ESI-MS of PC species in SM43Δ*cdsA* (D), and (E) the ESI-MS of PC species in SM43Δ*pgsA*. The mass spectra shown are averaged from spectra acquired by NPLC-ESI/MS during the 20–21 min window. PC species are detected by positive ion ESI/MS as the M^+^ ions.

### SM43 encodes a partial CDP-choline pathway

The CDP-choline pathway requires activation of choline to CDP-choline by the LicA and LicC enzymes. This is followed by condensation of CDP-choline with DAG by a 1,2-diacylglycerol choline phosphotransferase (CPT) enzyme to form PC (Fig 1). *S. mitis* and *S. oralis* encode *licABC* for the activation of exogenous choline to CDP-choline, which is required for choline decoration of Type IV LTA in these organisms and *S. pneumoniae* (38–42). Given that SM43 encodes *licABC*, we assessed the possibility that the CDP-choline pathway is used for PC biosynthesis in this organism.

The CPT of *T. denticola* possesses a CDP-alcohol phosphatidyltransferase domain (Superfamily accession: cl00453) (37). Only one SM43 predicted protein, phosphatidylglycerophosphate synthase (PgsA), possesses this domain. PgsA catalyzes the addition of glycerolphosphate to CDP-DAG to form phosphatidylglycerophosphate (PG-P) (55) which is required for subsequent PG and CL synthesis (Fig 1).

To investigate if PgsA has CPT activity in SM43, a 284 bp region of *pgsA* encoding the catalytic domain was replaced with an erythromycin resistance cassette. The *pgsA* mutant has a significant growth defect, with a doubling time almost twice that of the wild type (Fig S2a). The defect is likely due to PgsA interaction with RodZ in membrane homeostasis, as has been reported for *Bacillus subtilis* (56). Similar to SM43Δ*cdsA*, PC is present in SM43Δ*pgsA* cultured in THB (Fig 2e). As expected, the SM43Δ*pgsA* mutant lacks PG and CL (Table 1) and is high-level daptomycin-resistant (MIC >256 µg/mL), in agreement with experiments by Tran *et al*. utilizing *S. mitis-oralis* strains (57). We conclude that the sole candidate for CPT activity in SM43, PgsA, does not catalyze PC biosynthesis.

To conclusively exclude the CDP-choline pathway, we performed stable isotope-labeling experiments. SM43 was cultured in THB supplemented with 2 mM deuterated choline (choline-*d9*), in which all nine hydrogen (1 Da) atoms on the three methyl groups are replaced with deuterium (2 Da) atoms, thereby increasing the mass of choline by 9 Da. If SM43 utilizes the CDP-choline pathway for PC biosynthesis, an *m/z* shift of 9 Da would be observed for CDP-choline and PC species in choline-*d9*-supplemented SM43 cultures. We observed choline-*d9* incorporation into CDP-choline via a shift of its M^+^ ion from *m/z* 489 (Fig 3a; no choline-*d9* supplementation) to *m/z* 498 (Fig 3b, with choline-*d9* supplementation). The identification of CDP-choline and CDP-choline-*d9* was confirmed by MS/MS (Fig S3). In contrast, no *m/z* shift was observed for PC species between control (Fig 3c) and choline-*d9*-supplemented (Fig 3d) cultures. These data definitively eliminate the CDP-choline pathway as the SM43 PC biosynthesis pathway.

**Fig 3.**
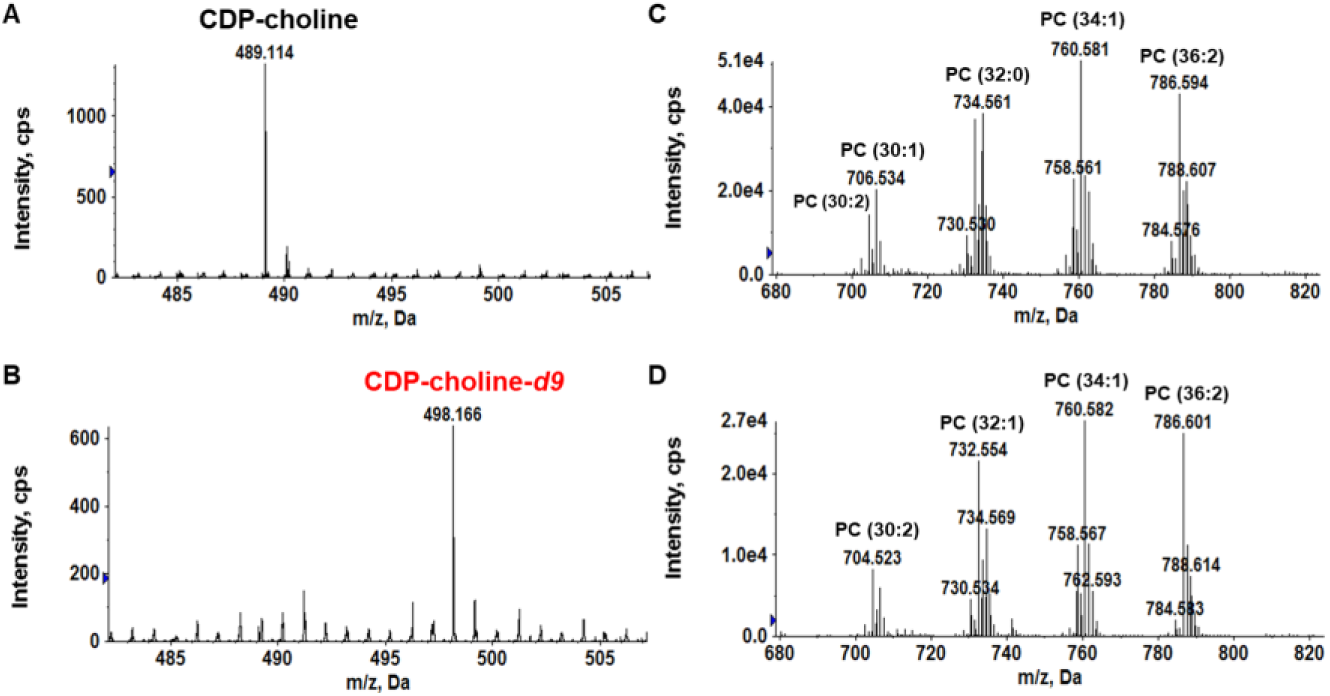
Exogenous deuterated choline (choline-*d9*) is used by SM43 to synthesize CDP-choline, but not PC. CDP-choline and PC were detected in the soluble-metabolite extract and lipid extract, respectively. Soluble metabolites were analyzed by reverse phase LC/ESI-MS in positive ion mode. Lipids were analyzed by normal phase LC/ESI-MS in positive ion mode. Shown are CDP-choline (M^+^ ion at *m/z* 489.1) present in SM43 cultured in THB (A), and CDP-choline-*d9* (M^+^ ion at *m/z* 498.1) present in SM43 cultured in THB supplemented with 2 mM choline-*d9* (B). Note the expected mass shift (9 Da) between CDP-choline and CDP-choline-*d9.* Also shown are PC species detected in SM43 cultured in THB (C), and PC species detected in SM43 cultured in THB supplemented with choline-*d9* (D). No corresponding mass shift (9 Da) is detected in the PC species, excluding the possibility of SM43 using the CDP-choline pathway for PC synthesis.

### SM43 utilizes the GPC scavenge pathway for PC biosynthesis

GPC is a major human metabolite, present in blood and saliva at concentrations up to 40 µM and 10 µM respectively (46). Using reverse-phase LC/MS, we detected GPC (*m/z* 258.1) in THB (Fig S4), likely originating from the heart infusion component of the medium. Therefore, GPC is available for scavenge in the medium used for routine SM43 cultures.

To determine whether SM43 utilizes the GPC pathway for PC biosynthesis, we used stable isotope-labeled GPC to trace the conversion of GPC into PC. SM43 was cultured in THB with and without 0.13 mM GPC-*d9* supplementation. A *m/z* shift of 9 Da was observed for all PC species (Fig 4a), demonstrating that SM43 uses the GPC pathway for PC biosynthesis. To further confirm this result, SM43 and *S. mitis* ATCC 49456 were cultured in a chemically defined medium containing 0.5 mM choline (DM) (58, 59), with and without 0.13 mM GPC supplementation. SM43 and ATCC 49456 synthesize PC only when GPC is present in the defined medium (Fig 5 and Table 2). In summary, GPC-*d9* isotope tracking and defined medium experiments independently confirm that SM43 utilizes the GPC pathway for PC biosynthesis. Moreover, these results are not strain-specific, as the *S. mitis* type strain also synthesizes PC only when GPC is present in the growth environment.

**Fig 4.**
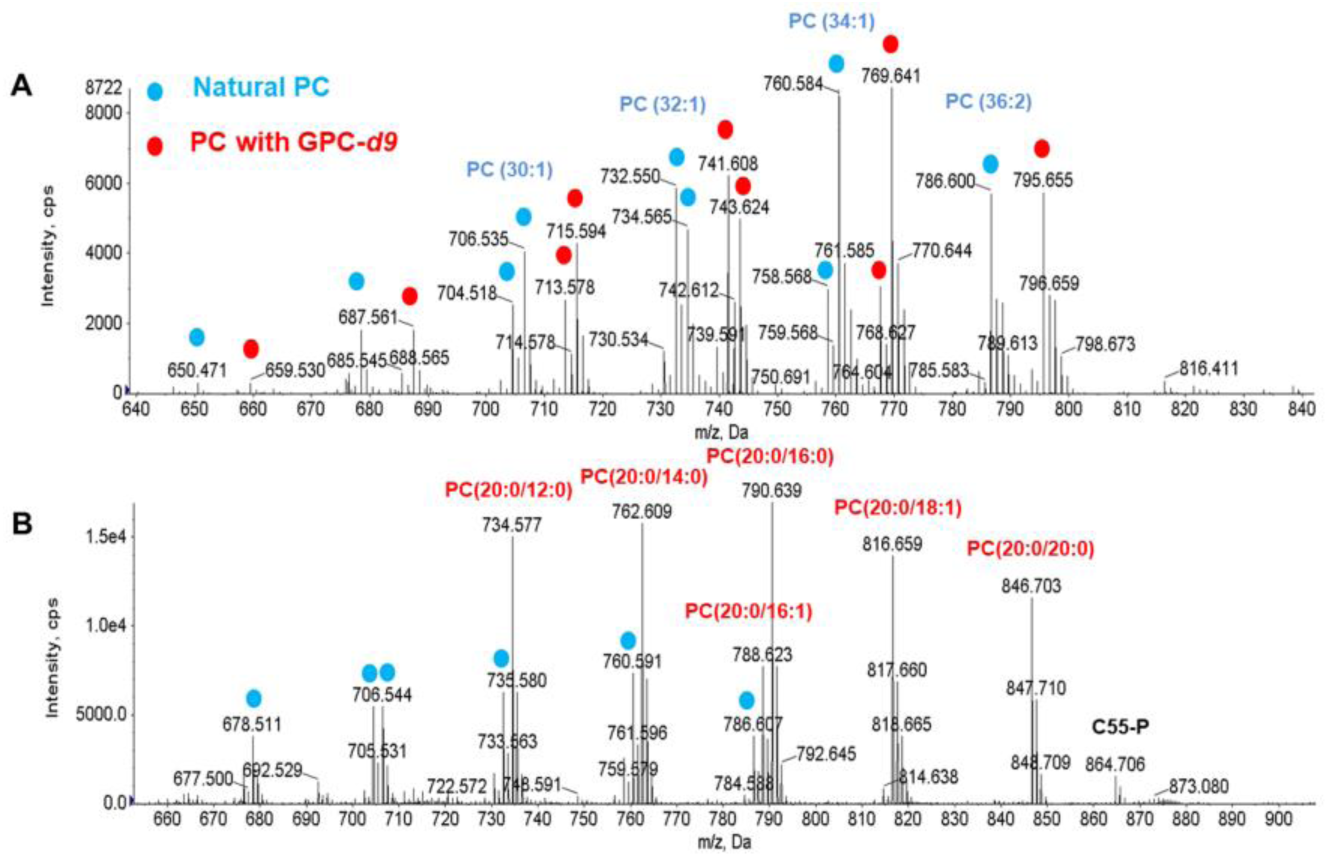
PC species of SM43 cultured in THB supplemented with GPC-*d9* and lysoPC (20:0). A) ESI/MS detection of PC species in SM43 cultured with GPC-*d9*. Blue dots indicate PC species normally detected in THB-grown SM43, and red dots indicate GPC-*d9*-originating PC species. B) ESI/MS detection of PC species in SM43 when cultured in the presence of lysoPC (20:0). Blue dots indicate PC species normally detected in THB-grown SM43. Red dots are lysoPC (20:0)-originating species. Incorporation of GPC-*d9* and lysoPC (20:0) into PC indicates that the GPC pathway is utilized by SM43 for PC biosynthesis.

**Fig 5.**
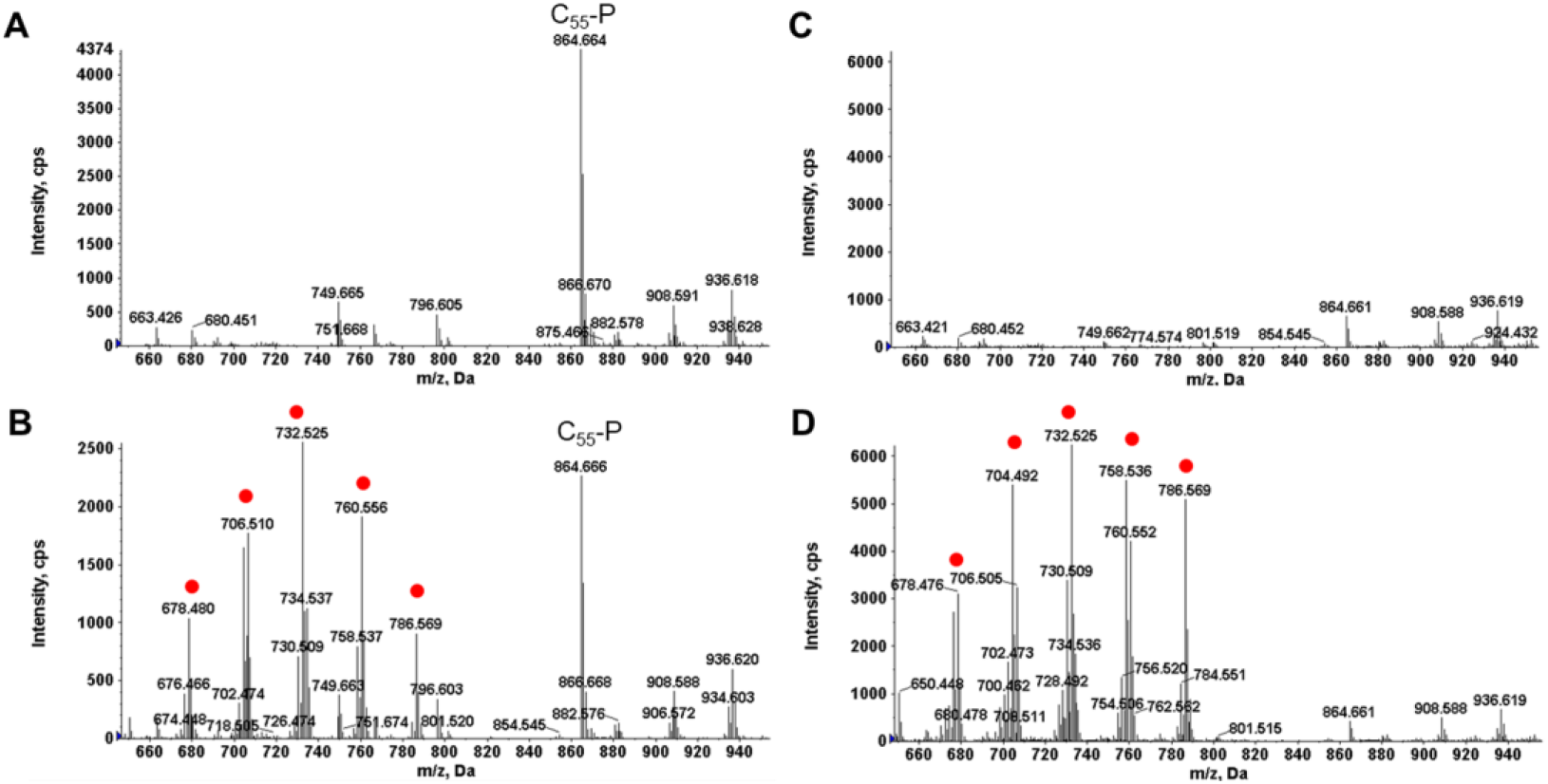
PC species of SM43 and *S. mitis* ATCC 49456 cultured in defined media (DM) with and without GPC. PC species (red dots) detected in SM43 (A) and ATCC 49456 (C) cultured in DM, and SM43 (B) and ATCC 49456 (D) cultured in DM supplemented with GPC are shown. PC is only detected when GPC is present in the culture medium.

**Table 2.** Summary of major glycolipids and phospholipids detected in Mitis group streptococci cultured in DM with or without GPC^1^.

| Streptococcal species | Strain | Source | DAG | MHDAG | DHDAG | PA | PG | CL | C55-P | PC | L-PG | Growth medium<sup>2</sup> |
| --- | --- | --- | --- | --- | --- | --- | --- | --- | --- | --- | --- | --- |
| *S. mitis* | ATCC 49456 | Type strain | + | + | + | + | + | + | + | - | - | DM |
|  | ATCC 49456 |  | + | + | + | + | + | + | + | + | - | DM+GPC |
| *S. oralis* | ATCC 35037 | Type strain | + | + | + | + | + | + | + | - | - | DM |
|  | ATCC 35037 |  | + | + | + | + | + | + | + | + | - | DM+GPC |
| *S. pneumoniae* | D39 | Historical strain | + | + | + | + | + | + | + | - | - | DM |
|  | D39 |  | + | + | + | + | + | + | + | + | - | DM+GPC |
| Mitis group | 1643 (SM43) | Endocarditis | + | + | + | + | + | + | + | - | - | DM |
|  | 1643 (SM43) |  | + | + | + | + | + | + | + | + | - | DM+GPC |

<sup>1</sup>Abbreviations: DAG, diacylglycerol; MHDAG, monohexosyldiacylglycerol; DHDAG, dihexosyldiacylglycerol; PA, phosphatidic acid; PG, phosphatidylglycerol; CL, cardiolipin; C55-P, undecaprenyl phosphate; PC, phosphatidylcholine; L-PG, lysyl-phosphatidylglycerol
<sup>2</sup>Defined media (DM); Defined media supplemented with glycerophosphocholine (DM+GPC).

PC is not an essential component in the lipid membrane for SM43 and ATCC 49456, evidenced by their abilities to grow in DM lacking GPC. To assess the impact that PC has on growth dynamics, SM43 and ATCC 49456 were cultured in THB and in DM with and without GPC supplementation. GPC presence or absence had no impact on growth (Fig S2b,c).

Since lysoPC is an intermediate in the GPC pathway, we hypothesized that SM43 could also scavenge lysoPC from its environment. LysoPC is present in human blood at up to 200 µM (60). We supplemented THB with lysoPC (20:0), which has an acyl chain that is not usually observed in bacterial membranes. SM43 readily scavenged exogenous lysoPC (20:0) and acylated it to form PC (Fig 4b). The PG species in the SM43 membrane remain unchanged in the presence of lysoPC (20:0), indicating low transacylation activity (Fig S5). We conclude that SM43 scavenges both GPC and lysoPC from its environment to synthesize PC (Fig 6).

**Fig 6.**
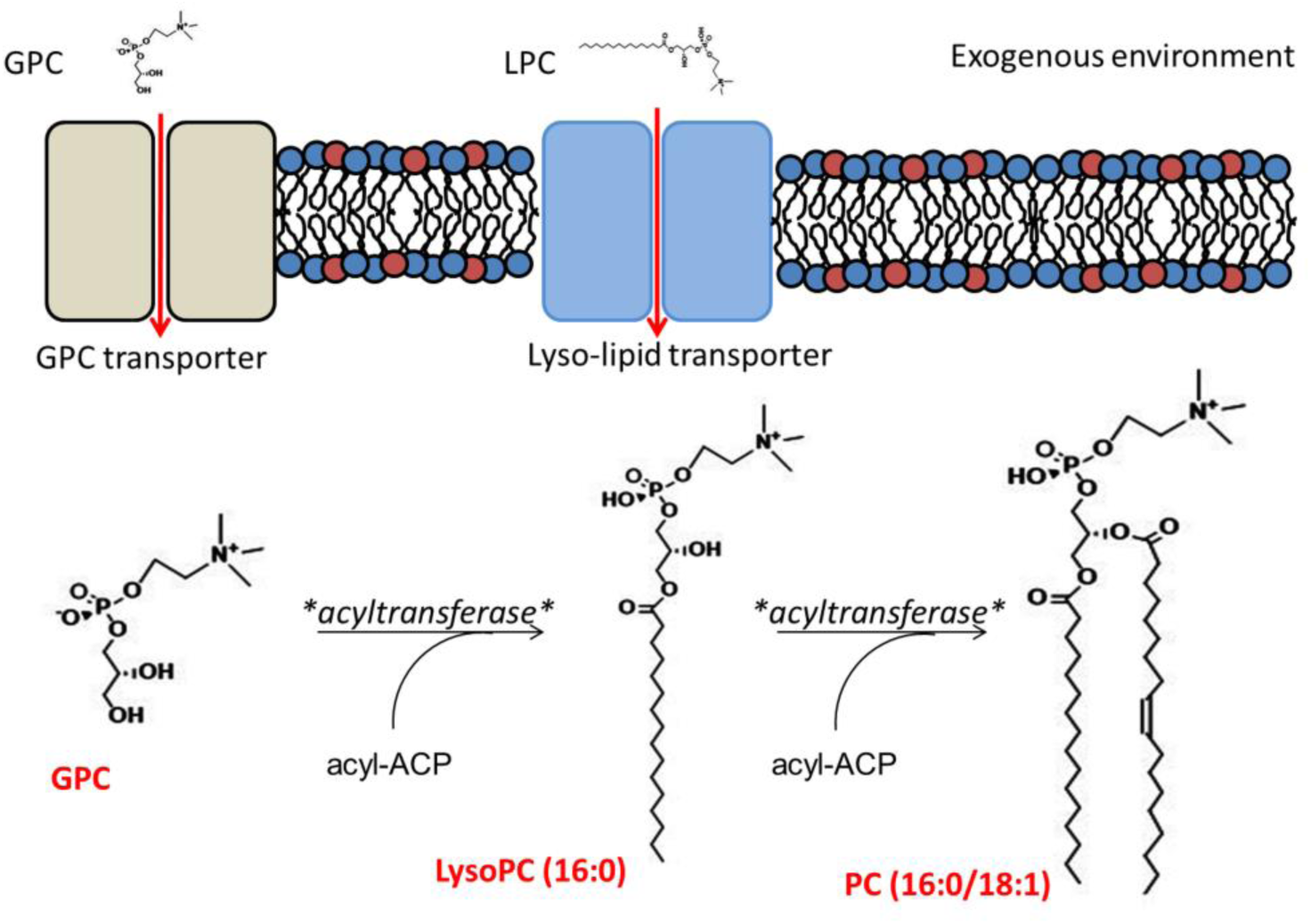
Proposed model for the GPC pathway in Mitis group streptococci. Exogenous GPC and lysoPC are transported into the cell via unidentified transporters. GPC is sequentially acylated to form lysoPC and PC. Acyl chain lengths vary in the lipid membrane; representative chain lengths are shown.

### GPC-dependent PC synthesis by *S. pneumoniae and S. oralis*

*S. pneumoniae* is a major human pathogen and a close relative of *S. mitis*. Surprisingly, there are only a few reports on lipid analysis of *S. pneumoniae* (17–19), for which thin layer chromatography was used as the analytical technique. We applied LC-ESI/MS, with much higher sensitivity and specificity, to investigate the lipidome of *S. pneumoniae*. PC is present in the membranes of *S. pneumoniae* D39 (Fig 7a,b) and TIGR4 (Fig 7c,d) cultured in THB supplemented with yeast extract (also see Table 1). Fig 7a and c shows the positive ion TIC and the relative abundance of PC in the D39 and TIGR4, respectively, in early stationary phase. The ESI/MS spectra of the major PC species, PC (30:1) at *m/z* 704, PC (32:1) at *m/z* 732, PC (34:2) at *m/z* 758, and PC(36:2) at *m/z* 786 are shown for D39 (Fig 7b) and TIGR4 (Fig 7d). We identified PC presence in infective endocarditis *S. oralis* isolates 1647 and 1648 in a previous study (14).

**Fig 7.**
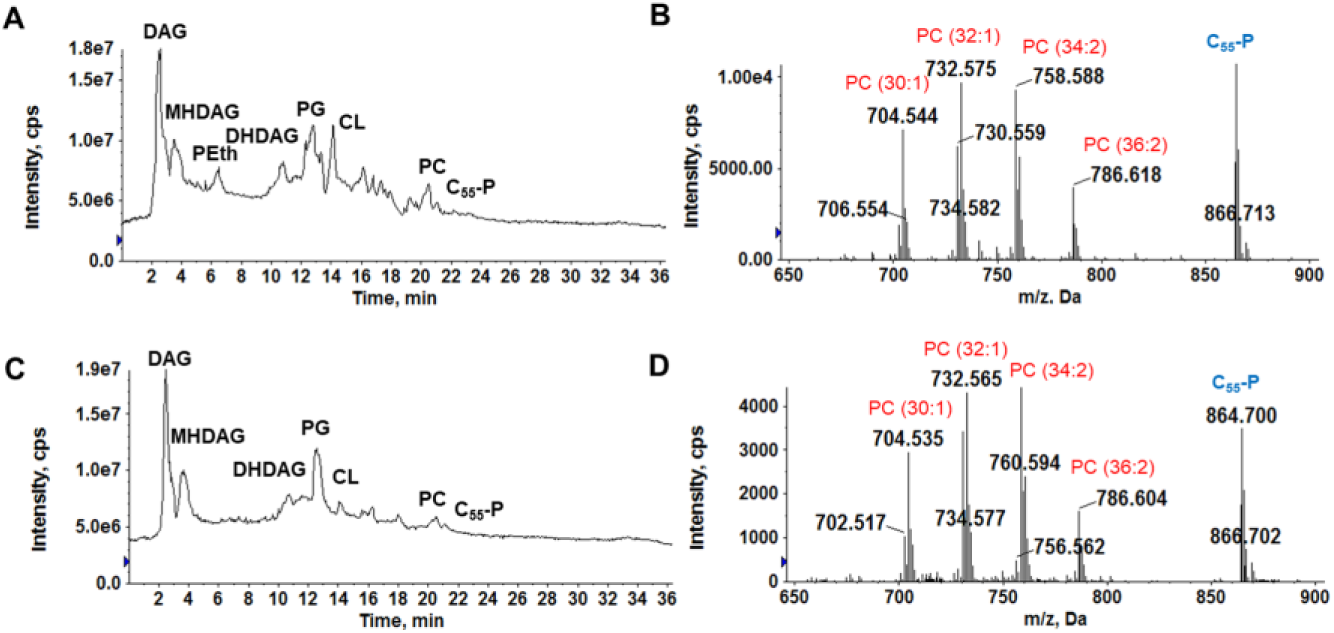
Positive ion total ion chromatogram (TIC) and PC mass spectrum of *S. pneumoniae* D39 and TIGR4 strains in early stationary phase. Shown are the TIC of *S. pneumoniae* D39 lipids (A), the ESI/MS of major PC species in *S. pneumoniae* D39 (B), the TIC of *S. pneumoniae* TIGR4 lipids (C), and the ESI/MS of major PC species in *S. pneumoniae* TIGR4 (D). The mass spectra shown are averaged from spectra acquired by NPLC-ESI/MS during the 19.5 – 21 min window. PC species are detected by positive ion ESI/MS as the M^+^ ions, while the co-eluting undecaprenyl phosphate (C_55_-P) is detected as the [M+NH_4_]^+^ ion (*m/z* 864.7).

To determine if *S. pneumoniae* and *S. oralis* utilize the GPC pathway for PC biosynthesis, *S. pneumoniae* D39 and *S. oralis* ATCC 35037 were cultured in defined media with and without GPC supplementation. When GPC is not available, *S. pneumoniae* and *S. oralis* do not synthesize PC (Fig 8a,c). PC is only present when GPC is available in the medium (Fig 8b,d). We conclude that *S. pneumoniae* and *S. oralis* also scavenge GPC to synthesize PC.

**Fig 8.**
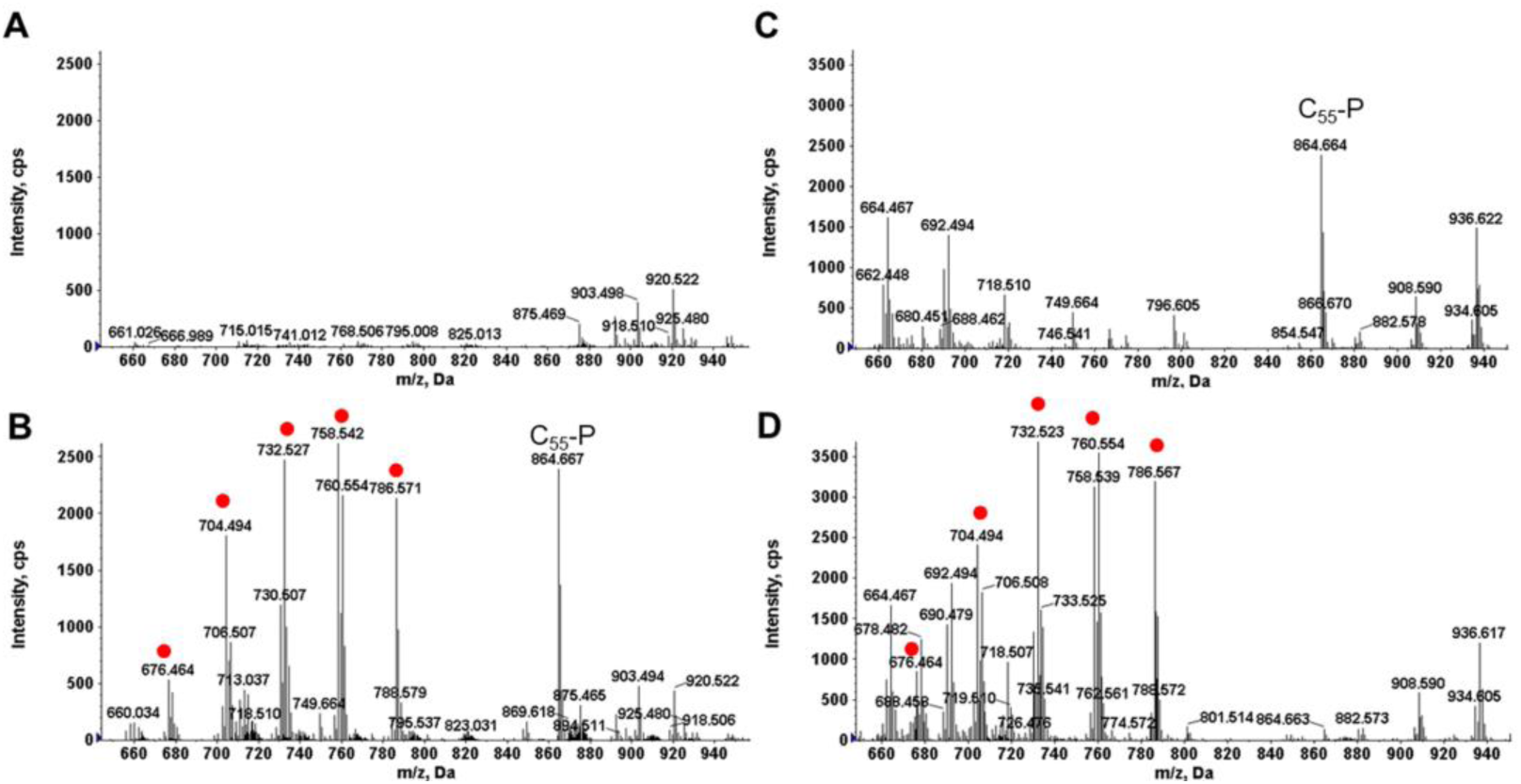
GPC pathway confirmation in *S. pneumoniae* D39 and *S. oralis* ATCC 35037. Shown are PC species (red dots) detected for D39 (A) and ATCC 35037 (C) cultured in DM and D39 (B) and ATCC 35037 (D) cultured in DM supplemented with GPC. PC is only detected when GPC is present in the culture medium.

## Discussion

The lipid membrane is a dynamic site of interaction between microbial pathogens and their hosts. However, for many pathogens, the composition of the membrane is poorly understood. In this study, we characterized the lipidomes of selected species of Mitis group streptococci and investigated the mechanistic basis for biosynthesis of the phospholipid PC. We found that Mitis group streptococci remodel their membrane lipid compositions in response to the host metabolites GPC and lysoPC. To our knowledge, this is the first description of PC in *S. pneumoniae*, a major human pathogen which has been studied for over a century and yet its membrane lipid composition remains poorly understood. There have been very few lipidomic studies performed in *S. pneumoniae* (17–19), and little is known about how *S. pneumoniae* remodels its membrane in response to changing environments inside and outside the host. Here, we reported the first identification of PC in *S. pneumoniae* and demonstrated that *S. pneumoniae* synthesizes PC only when GPC, a major human metabolite, is available for scavenge.

Very little is known about the GPC in bacteria; however, a complete GPC pathway has been characterized in yeast (44, 48). It includes a dual substrate transporter, Git1, for the uptake of glycerophosphoinositol and GPC; a GPC acyltransferase referred to as Gpc1; and the acyltransferase Ale1 performing lysoPC acylation. To fully elucidate the GPC pathway in streptococci, identification of transporters for GPC and lysoPC as well as the acyltransferases are required. However, no orthologs of the yeast GPC pathway components were identified in the SM43 genome. There are two acyltransferases annotated in the *S. mitis* genome, PlsY and PlsC, which are responsible for phosphatidic acid biosynthesis in other organisms and may play a role in acylation of either GPC or lysoPC. In *Mycobacterium tuberculosis*, a sugar-binding ABC transporter, UgpABCE, transports GPC (61). The UgpB substrate binding domain is flexible in substrate affinity, binding to phosphate-containing substrates such as *sn*-glycerol 3-phopshate, glycerol 2-phosphate, and GPC (61–63). Given the reduced size of streptococcal genomes, it is possible that GPC uptake is also conferred by a transport system with flexible substrate specificity.

Is PC biosynthesis important for Mitis group streptococcal virulence? We expect that PC levels in the membrane will impact membrane charge, for example, which could in turn impact biofilm formation and interactions with the host immune system. Moreover, GPC and lysoPC levels vary at different sites within the human body and in health and disease (≤ 500 µM and ≤ 200 µM, respectively (46, 60)), which could impact the relative ratio of the zwitterionic PC to the anionic phospholipids PG and CL in Mitis group streptococci colonizing these sites. Due to our limited understanding of the genes underlying the GPC pathway, at present our only method to control this pathway is by altering the *in vitro* growth medium. For this reason *in vivo* studies are not currently feasible. However, by culturing Mitis group streptococci in defined media with and without GPC, in future studies we can assess the biophysical impact of PC on streptococcal membranes in terms of lipid/microdomain organization, charge, rigidity, and protein composition, which would be informative from a basic science perspective.

Overall, our work highlights the importance of utilizing laboratory culture media that mimic the *in vivo* nutritional environments where pathogens are found. Our identification of PC in the membrane of Mitis group streptococci including the major human pathogen *S. pneumoniae*, and their utilization of the rare GPC pathway, justifies further investigation into streptococcal membrane biology, about which little is known.

## Materials and methods

### Bacterial strains, media, and growth conditions

Strains and plasmids used in this study are shown in Table S1. Streptococcal strains were grown in THB in 5% CO_2_ at 37°C unless otherwise stated. *S. pneumoniae* THB cultures were supplemented with 0.5% yeast extract. Streptococcal chemically defined medium (58) was diluted from stock as described (59) supplemented with 0.5 mM choline (referred to as DM), slightly modified from (64) unless otherwise stated. Media were supplemented with 130 µM glycerophosphocholine (Sigma-Aldrich) where stated. Erythromycin was used at 20 µg/mL for SM43. Daptomycin susceptibilities were assessed using daptomycin Etest strips (bioMérieux) on Mueller Hinton Agar plates, per CLSI standards (65).

### Genome sequencing and assembly

Genomic DNA was isolated using the Qiagen DNeasy Blood and Tissue kit per the manufacturer’s protocol with the exception that cells were pre-treated with 180 µL 50 mg/mL lysozyme, 25 µL 2500 U/mL mutanolysin, and 15 µL 20 mg/mL pre-boiled RNase A and incubated at 37°C for 2 h. Pacific Biosciences single molecule real time (SMRT) sequencing was performed by the Johns Hopkins Genome Core. The SM43 whole genome was assembled using the Unicyler assembly pipeline (66) combining SMRT long reads generated in this study and Illumina reads we previously generated for SM43 (accession PRJNA354070; (14)). Sequencing of SM43Δ*pgsA* was performed by Molecular Research DNA (Shallowater, Texas) using Illumina Hi-Seq 2×150 paired end reads.

### Gene deletions in SM43

Primers used in this study are shown in Table S2. The SM43 *cdsA* deletion construct was designed essentially as previously described (67, 68). See Text S1. For *pgsA* deletion, the protocol was the same as for *cdsA* deletion, except an erythromycin resistance marker with its native promoter was amplified from pG^+^host4 (69) and inserted between the homologous flanking arms to allow for selection-based screening of putative *pgsA* mutants. The resulting construct deleted 284 bp encoding the catalytically active site of *pgsA* and replaced them with the erythromycin resistance marker. The replacement of *pgsA* was confirmed by whole genome sequencing.

### Natural transformation

The protocol was as previously described (68) with minor modifications. Briefly, 100 µL exponential phase pre-cultures (OD_600_ ∼0.5) in THB were frozen with an equal volume of 10% glycerol. Precultures were thawed at room temperature and diluted in 900 µL THB. This was further diluted 1:50 in pre-warmed 5 mL THB and incubated at 37°C for 45 minutes. 500 µL culture was aliquoted along with 1 µL of 1 mg/mL competence stimulating peptide (DWRISETIRNLIFPRRK) and 1 µg/mL linear DNA construct. Transformation reactions were cultured for 2 h at 37°C in microcentrifuge tubes before plating on THB agar with selection as appropriate.

### Lipidomics

Unless otherwise noted, lipidomics analyses were performed on overnight cultures in stationary phase. Centrifugation was performed using a Sorvall RC6+ centrifuge. Cultures were pelleted at 4,280 × *g* for 5 min at room temperature. The supernatants were removed and stored at −80°C until acidic Bligh-Dyer lipid extractions were performed as described (70) with minor modifications. See Text S1.

Normal-phase LC was performed on an Agilent 1200 quaternary LC system equipped with an Ascentis silica high-performance liquid chromatography (HPLC) column (5m; 25 cm by 2.1 mm; Sigma-Aldrich) as described previously (70, 71). Data analysis was performed using Analyst TF1.5 software (Sciex, Framingham, MA). See Text S1.

### Metabolite extractions

Cultures were pelleted at 4,280 × *g* in a Sorvall RC 6+ floor centrifuge at room temperature, washed once with 1X PBS, and transferred to 1.5 mL microcentrifuge tubes. Cells were pelleted and frozen at −80°C until use. Metabolite extraction was performed as previously described (72) with minor modifications. See Text S1.

### Reverse phase LC-ESI/MS analysis

RPLC-ESI/MS analysis of water-soluble metabolites was performed using a Shimadzu LC system (comprising a solvent degasser, two LC-10A pumps and a SCL-10A system controller) coupled to a TripleTOF5600 mass spectrometer (Sciex, Framingham, MA). See Text S1 for detailed information of LC flow rate and mass spectrometer settings. Data acquisition and analysis were performed using the Analyst TF1.5 software (Sciex, Framingham, MA).

### Deuterated isotope (*d9*) and lysoPC tracking

Deuterated (*d9*) isotope tracking was performed by addition of 2 mM choline-*d9* (Sigma-Aldrich) in 50 mL THB. 3.7 mM GPC-*d9* (Toronto Research Chemicals) was added to 15 mL THB, to yield a final concentration of 130 µM. Cultures were grown in 5% CO_2_ at 37°C for 18 h before 10 mL of choline-*d9* culture or 5 mL of GPC-*d9* culture was removed for metabolite extraction and the remaining culture was pelleted for lipid extraction. LysoPC (20:0) was obtained from Avanti Polar Lipids. Cultures were supplemented with 2 mg lysoPC per 15 mL THB, unless otherwise stated, and incubated overnight in 5% CO_2_ at 37°C. Lipid extractions were performed as described above.

### Growth curves

Individual colonies were incubated overnight in 5 mL DM. Cultures were diluted to a starting OD_600nm_ of 0.05 in pre-warmed 20 mL DM, DM with 130 µM GPC, and THB. OD_600nm_ was monitored every hour using a Thermo Scientific Genesys 30 spectrophotometer. Growth curves were performed in biological triplicates for each strain.

### ANI analysis

The ANI calculator (50, 51) was used with default parameters to analyze the following genomes: SM43, *S. oralis* ATCC 35037 (PRJNA38733) and *S. mitis* ATCC 49456 (PRJNA173).

### Accession numbers

The SM43 whole genome sequence generated in this study has been deposited in Genbank under the accession number CP040231. Illumina and SMRT sequence reads generated in this study have been deposited in the Sequence Read Archive under the accession number PRJNA542100.

## Acknowledgments

This work was supported by R01AI116610, R21AI130666, and the Cecil H. and Ida Green Chair in Systems Biology Science to K.P. and GM069338 and EY023666 to Z.G. We gratefully acknowledge Dr. Michael Federle and Jennifer Chang at the University of Illinois at Chicago for providing *S. pneumoniae* cell pellets from THB+Y for our original analysis and for hosting L.J. for *S. pneumoniae* culture work.

## Supplemental information

### Text S1: Supplementary Materials and Methods

#### Acidic Bligh-Dyer extractions

Centrifugation was performed using a Sorvall RC6+ centrifuge. Cultures were pelleted at 4,280 × *g* for 5 min at room temperature. The supernatants were removed and stored at −80°C until acidic Bligh-Dyer lipid extractions were performed as described (1) with minor modifications. Cell pellets were resuspended in 1X PBS (Sigma-Aldrich) and transferred to Coring Pyrex glass tubes with PTFR-lined caps (VWR), followed by 1:2 vol:vol chloroform:methanol addition. Single phase extractions were vortexed periodically and incubated at room temperature for 15 minutes before 500 × *g* centrifugation for 10 min. A two-phase Bligh-Dyer was achieved by addition of 100 µl 37% HCL, 1 mL CHCl_3_, and 900 µl of 1X PBS, which was then vortexed and centrifuged for 5 min at 500 × *g.* The lower phase was removed to a new tube and dried under nitrogen before being stored at −80°C prior to lipidomic analysis.

#### Normal phase Liquid Chromatography/ Mass Spectrometry

Normal-phase LC/MS was performed on an Agilent 1200 quaternary LC system equipped with an Ascentis silica high-performance liquid chromatography (HPLC) column (5m; 25 cm by 2.1 mm; Sigma-Aldrich) as described previously (1, 2). Briefly, mobile phase A consisted of chloroform-methanol-aqueous ammonium hydroxide (800:195:5, vol/vol), mobile phase B consisted of chloroform-methanol-water-aqueous ammonium hydroxide (600: 340:50:5, vol/vol), and mobile phase C consisted of chloroform-methanol-water-aqueous ammonium hydroxide (450:450:95:5, vol/vol/vol/vol). The elution program consisted of the following: 100% mobile phase A was held isocratically for 2 min, then linearly increased to 100% mobile phase B over 14 min, and held at 100% mobile phase B for 11 min. The LC gradient was then changed to 100% mobile phase C over 3 min, held at 100% mobile phase C for 3 min, and, finally, returned to 100% mobile phase A over 0.5 min and held at 100% mobile phase A for 5 min. The LC eluent (with a total flow rate of 300 ml/min) was introduced into the ESI source of a high-resolution TripleTOF5600 mass spectrometer (Sciex, Framingham, MA). Instrumental settings for negative-ion ESI and MS/MS analysis of lipid species were as follows: IS = −4,500 V, CUR = 20 psi, GSI = 20 psi, DP = −55 V, and FP = −150V. The MS/MS analysis used nitrogen as the collision gas. Data analysis was performed using Analyst TF1.5 software (Sciex, Framingham, MA).

#### Metabolite extractions

Briefly, cell pellets were resuspended in 300 µL super-chilled methanol:water (vol:vol 4:1) via vortexing and pipetting, and stored on dry ice for 15 min. Suspensions were centrifuged at 17,136 × *g* at 4°C for 5 min in a table top centrifuge. The supernatant was removed to a pre-chilled Corning Pyrex glass tube and kept on dry ice. A further 2 resuspensions in 200 µL methanol:water (vol:vol 4:1) were performed. Pooled supernatants were dried under nitrogen at room temperature and stored at −80°C prior to analysis.

#### Reverse phase LC-ESI/MS analysis

RPLC-ESI/MS analysis of water-soluble metabolites was performed using a Shimadzu LC system (comprising a solvent degasser, two LC-10A pumps and a SCL-10A system controller) coupled to a TripleTOF5600 mass spectrometer (Sciex, Framingham, MA). LC was operated at a flow rate of 200 μl/min with a linear gradient as follows: 100% of mobile phase A was held isocratically for 2 min and then linearly increased to 100% mobile phase B over 14 min and held at 100% B for 4 min. Mobile phase A consisted of methanol/acetonitrile/aqueous 1mM ammonium acetate (60/20/20, v/v/v). Mobile phase B consisted of 100% ethanol containing 1 mM ammonium acetate. A Zorbax SB-C8 reversed-phase column (5 μm, 2.1 × 50 mm) was obtained from Agilent (Palo Alto, CA). The LC eluent was introduced into the ESI source of the mass spectrometer. Instrument settings for negative ion ESI/MS and MS/MS analysis of lipid species were as follows: Ion spray voltage (IS) = −4500 V; Curtain gas (CUR) = 20 psi; Ion source gas 1 (GS1) = 20 psi; De-clustering potential (DP) = −55 V; Focusing potential (FP) = −150 V. Data acquisition and analysis were performed using the Analyst TF1.5 software (Sciex, Framingham, MA).

#### Deletion of SM43 *cdsA*

Primers used in this study are shown in Table S2. The *cdsA* deletion construct was designed essentially as previously described (3, 4). Approximately 2 Kb flanking regions on either side of the gene were amplified using Phusion polymerase (Thermo Fisher). PCR products were digested with restriction enzymes (New England Biolabs) and ligated using T4 DNA ligase (New England Biolabs). Ligations were amplified by PCR utilizing the 5’ and 3’ most primers with Phusion polymerase to obtain a linear product. The PCR product was analyzed on a 1% agarose gel and extracted using the QIAquick Gel Extraction Kit per the manufacturer’s protocol. The linear construct was transformed into SM43 via natural transformation, described below. Transformation plates were incubated overnight. Putative transformant colonies were inoculated into THB and screened via PCR for the clean deletion of *cdsA*. The *cdsA* regions of putative mutants were sequenced for confirmation (Massachusetts General Hospital DNA Core).

**Fig S1.**
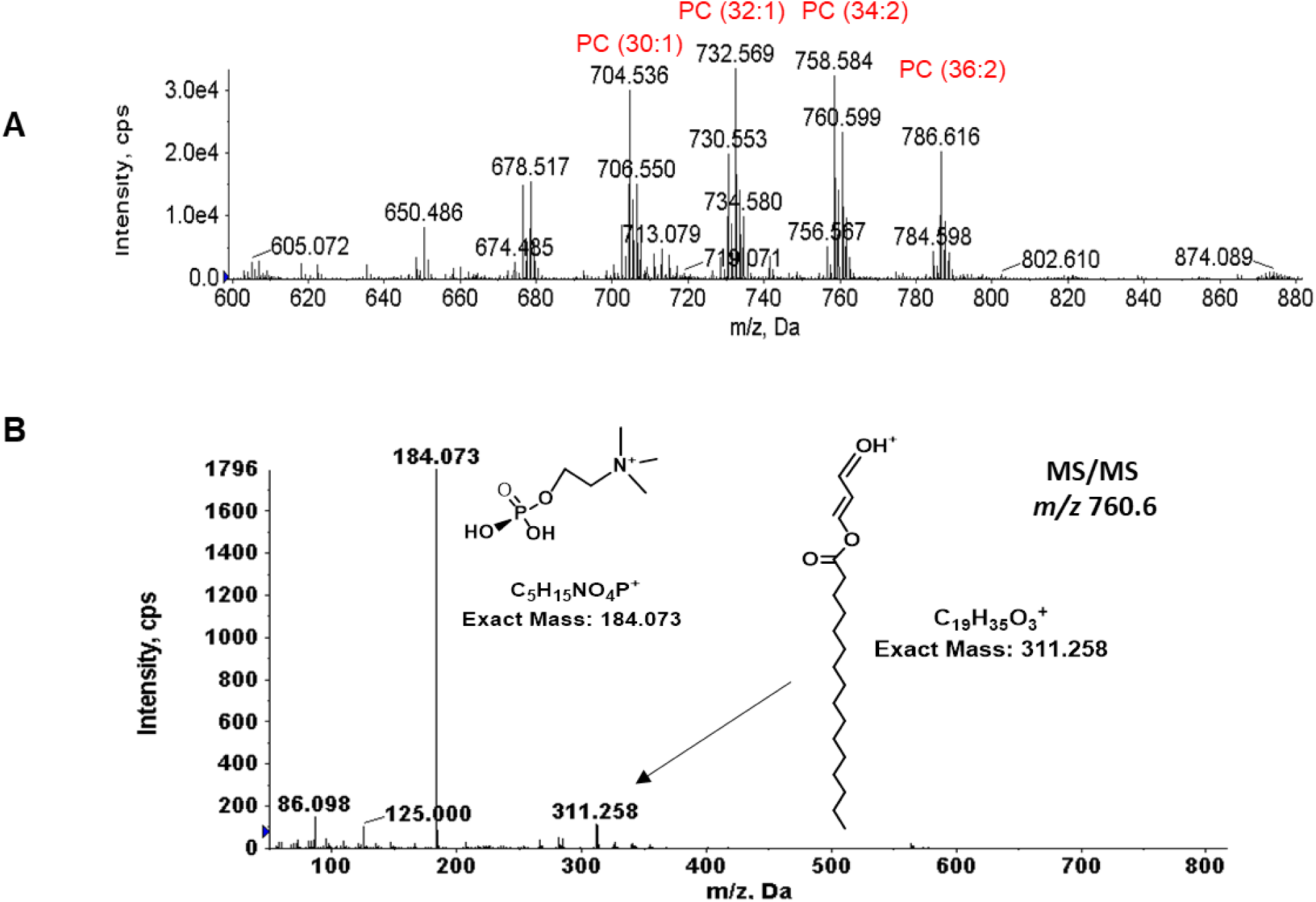
Phosphatidylcholine (PC) species in *S. mitis* ATCC 49456. A) *S. mitis* ATCC 49456, B) MS/MS daughter ion fragmentation of PC (34:1) *m/z* 760.6 in SM43. The mass spectra shown are averaged from spectra acquired by NPLC-ESI/MS during the 20–21 min window. PC species are detected by positive ion ESI/MS as the M^+^ ions.

**Fig S2.**
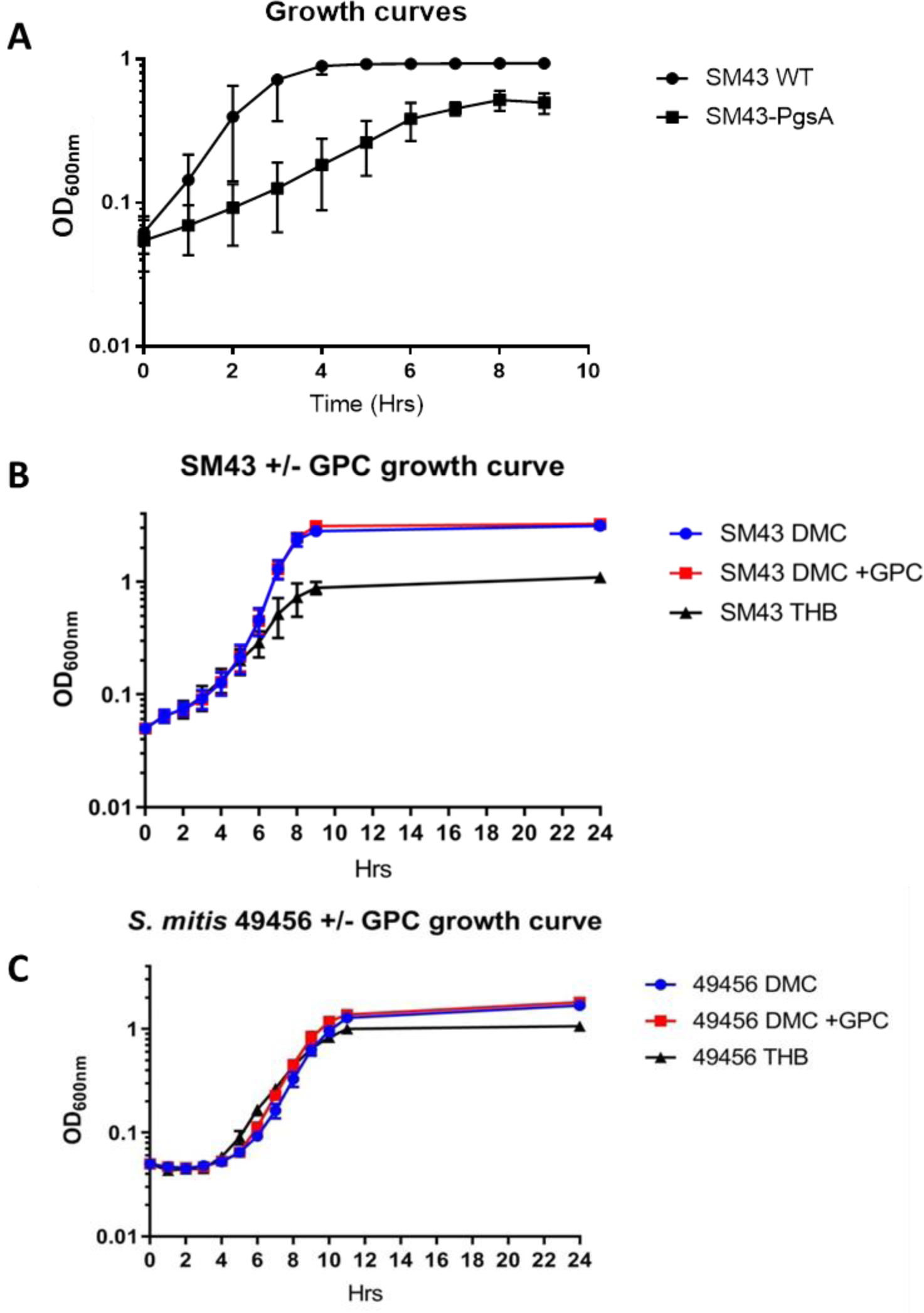
Growth curves. A) Growth curves for SM43 and SM43Δ*pgsA* in THB. B) Growth curves of SM43 in THB, and in defined medium (DM) with and without GPC supplementation and C) Growth curves of ATCC49456 in THB, and in defined medium (DM) with and without GPC supplementation. OD_600nm_ readings were taken every hour, with a starting inoculum of OD_600nm_ 0.05. Growth curves were performed as biological triplicate. Error bars indicate standard deviation.

**Fig S3.**
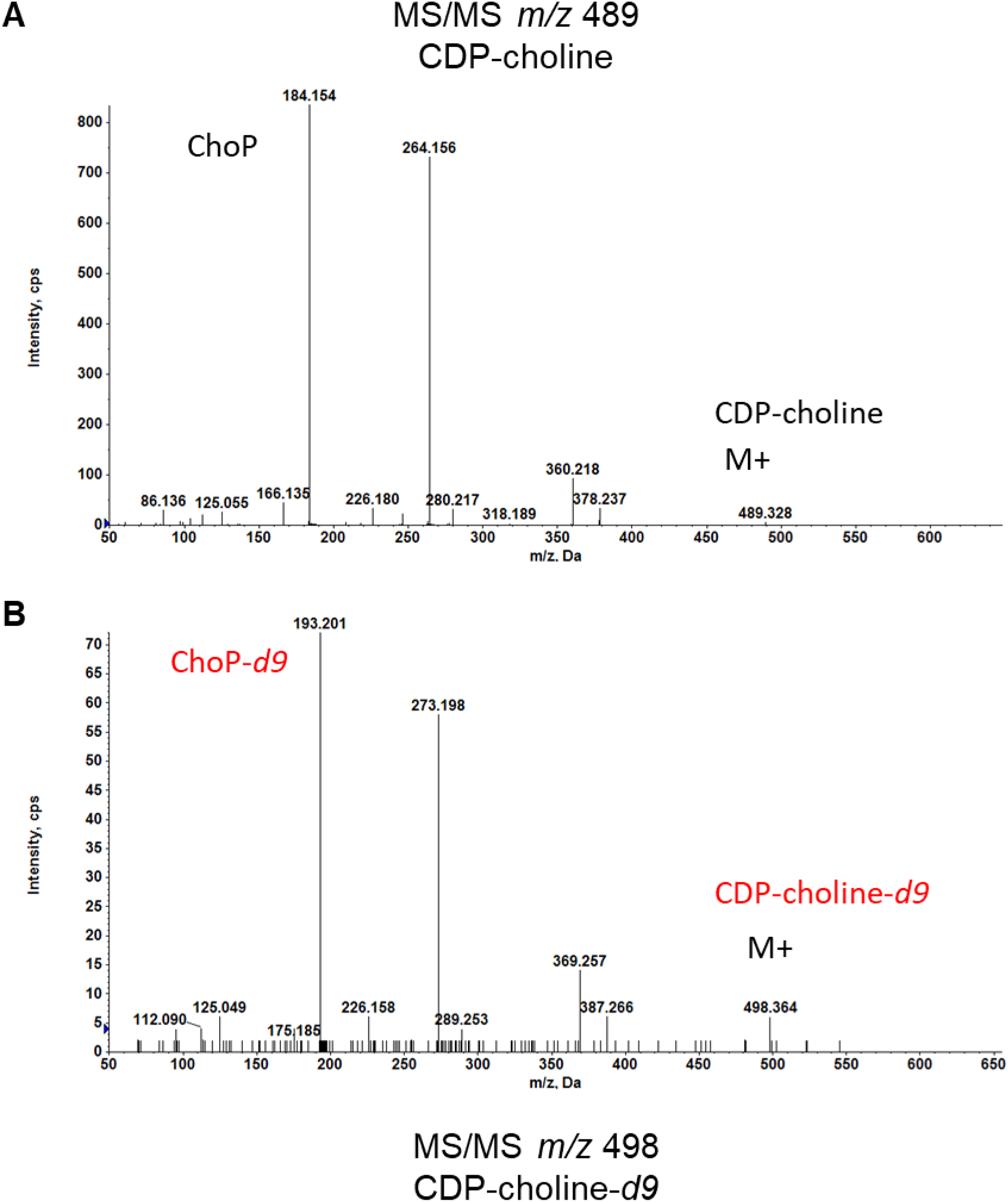
MS/MS fragmentation of CDP-choline and CDP-choline-*d9*. A) MS/MS of CDP-choline (*m/z* 489) B) MS/MS of CDP-choline-*d9* (*m/z* 498). Note the mass shift of 9-*da* for all major MS/MS peaks in B.

**Fig S4.**
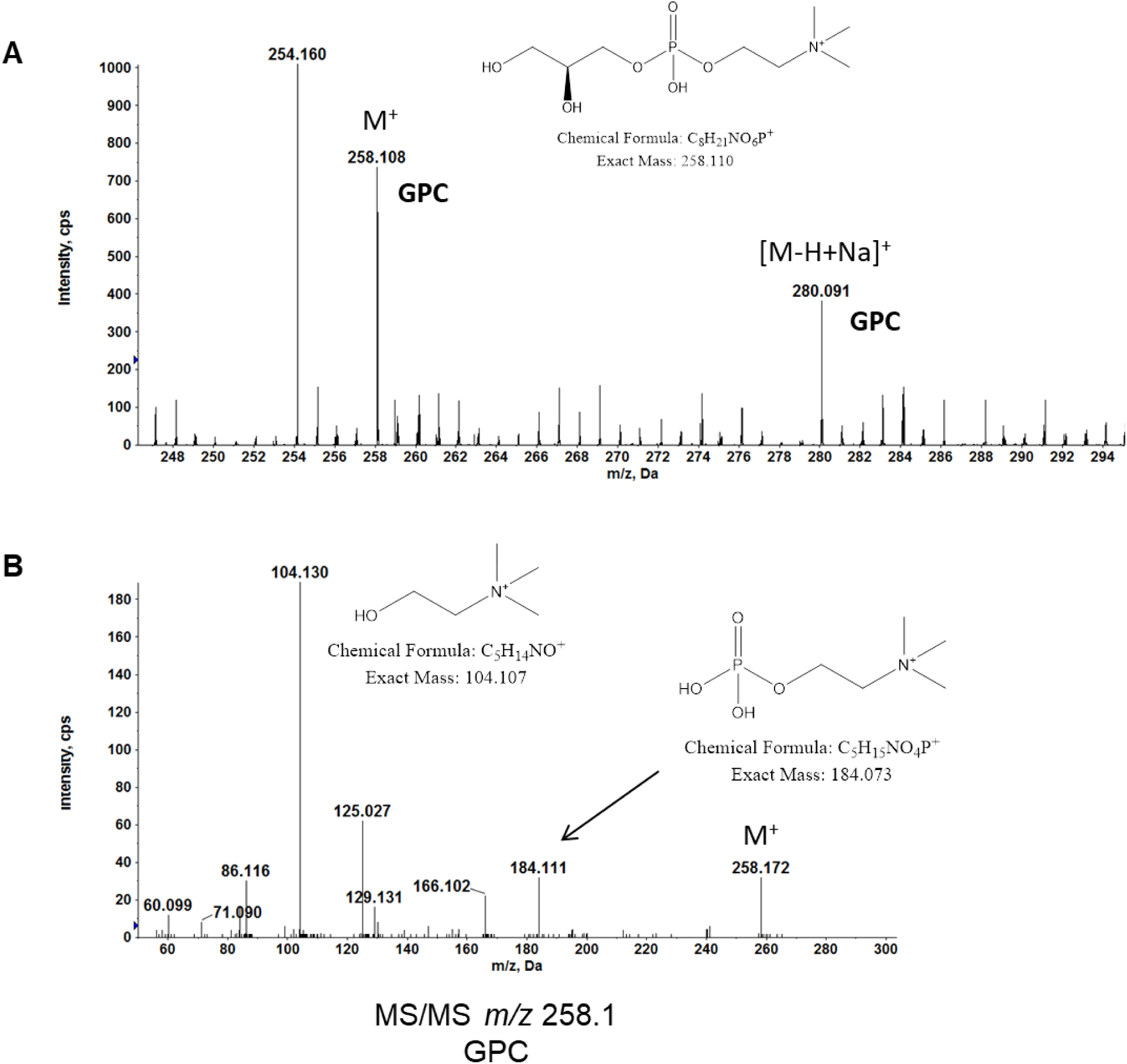
MS detection of GPC in THB. The soluble-metabolite extract of THB was analyzed by reverse phase LC/MS. A) GPC is detected by positive ion ESI/MS as the M^+^ ion at *m/z* 258.108 and [M-H+Na]^+^ ion at *m/z* 280.091. B) MS/MS spectrum of the M^+^ ion (*m/z* 258.1) of GPC. The chemical structures of major fragments are shown.

**Fig S5.**
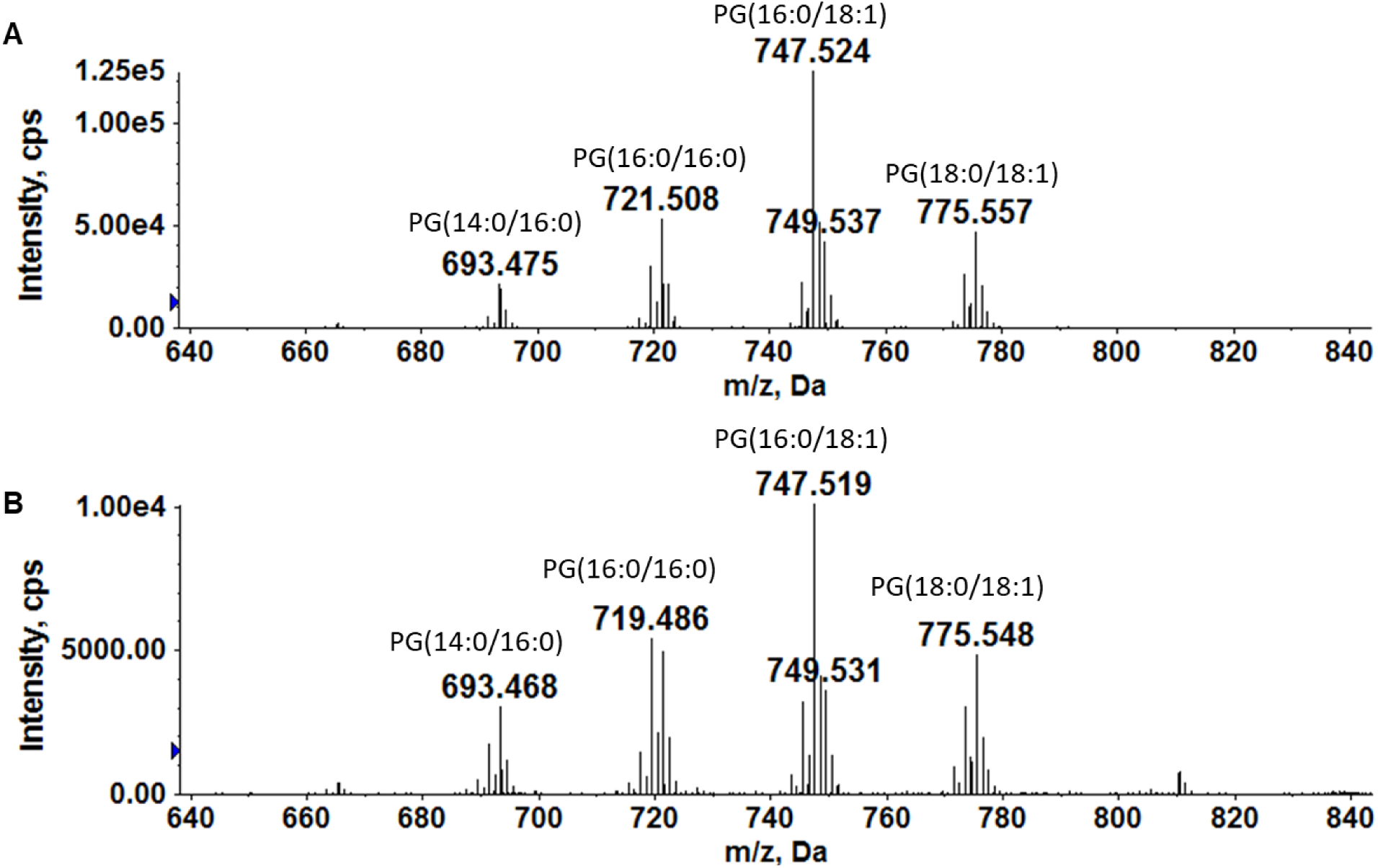
ESI/MS of phosphatidylglycerol (PG) species of SM43 cultured in THB with and without lysoPC (20:0) addition. A) ESI/MS detection of PG species in SM43 cultured in THB. B) ESI/MS detection of PG species in SM43 cultured in THB in the presence of lysoPC (20:0). In sharp contrast to what is observed for PC, there is no appreciable incorporation of the 20:0-acyl chain into PG in the presence of lysoPC (20:0). The lack of incorporation of C20:0 into PG indicates the insignificant transacylation activity of SM43. PG species are detected by negative ion ESI/MS as the [M-H]^−^ ions.

**Table S1.** Strains and plasmids used in this study.

| Organism | Strain | Description | Ref |
| --- | --- | --- | --- |
| Mitis group streptococci | 1643 (SM43) | Wild-type infective endocarditis isolate | (5) |
| | 1643Δ*cdsA* | *cdsA* deletion strain | This work |
| | 1643Δ*pgsA::ermB* | *pgsA* deletion strain, replaced with *ermB*. Erm<sup>r</sup> | This work |
| *S. mitis* | ATCC 49456 (NCTC 12261) | Wild-type *S. mitis* type strain from ATCC | (6) |
| *S. oralis* | ATCC 35037 (NCTC 11427) | Wild-type *S. oralis* type strain from ATCC | (7) |
| *S. pneumoniae* | D39 | Wild type, historical strain | (8) |
|  | TIGR4 | Wild type, bloodstream isolate | (9) |
| Plasmid | Description |  | Ref |
| pG<sup>+</sup>host 4 | Encodes *ermB* |  | (10) |

**Table S2.**
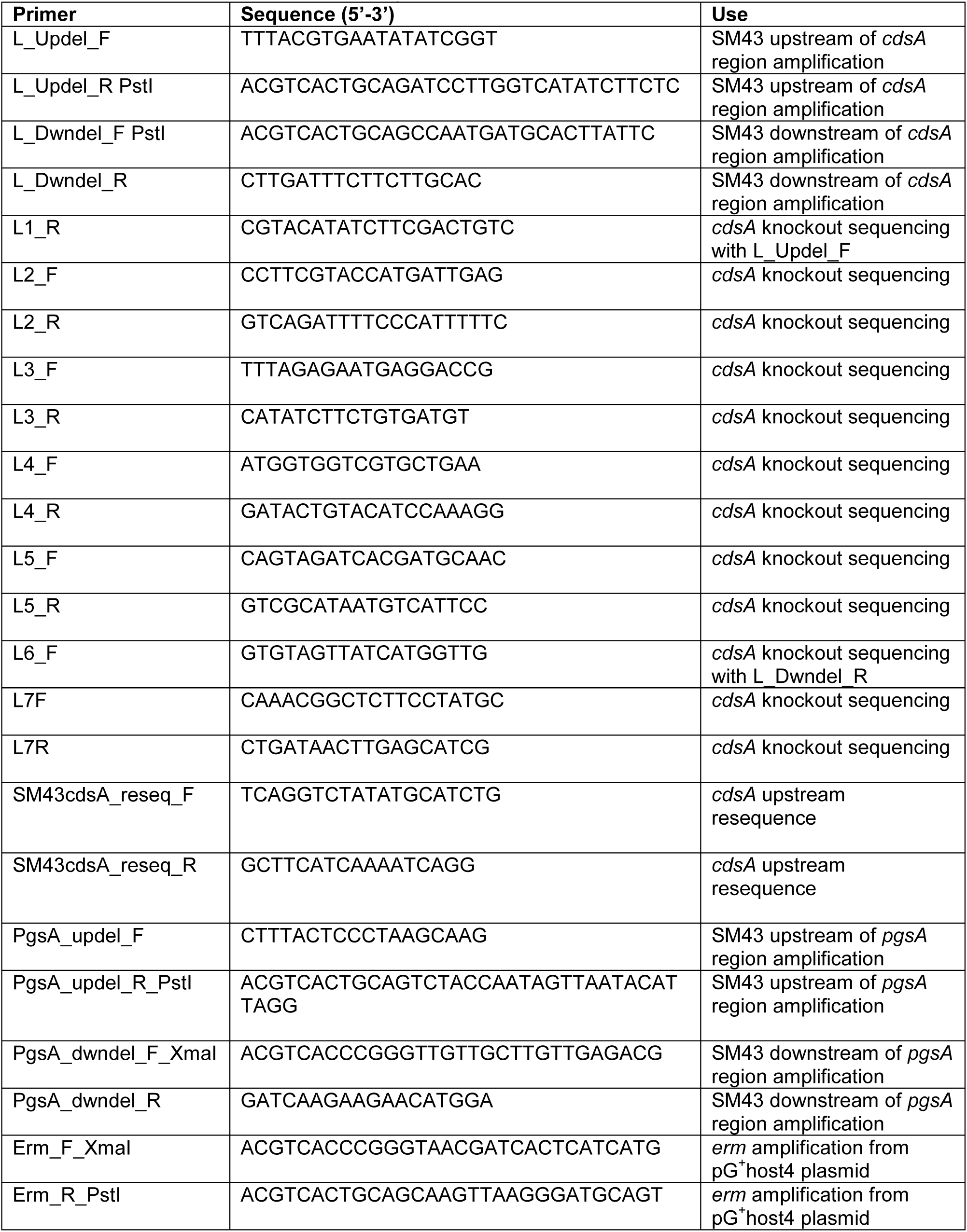
Primers used in this study.

